# Using large language models for enhancing accessibility for Monte Carlo photon transport simulations and beyond

**DOI:** 10.64898/2026.07.20.738933

**Authors:** Fan-Yu Yen, Yiyi Liu, Qianqian Fang

## Abstract

**Significance:** Computational modeling and the use of simulation software tools are essential for biomedical optics research. Designing effective simulations often requires in-depth understanding of the underlying physical problems and proper configuration of the software settings, which often constitute key barriers for novice users including students. The rapid emergence of large language models (LLMs) offers new opportunities for natural-language-based interaction, but integrating them with technical software remains challenging because of their limited output reproducibility. Overcoming these limitations would allow more intuitive, efficient, and reproducible interaction between scientists and scientific software.

**Aim:** We investigate the use of LLMs in quantitative biophotonics simulation tools, with a goal of enabling novice users to build complex photon simulations using intuitive natural-language-based problem descriptions.

**Approach:** We have explored prompt engineering strategies that enable LLMs to bridge the gap between natural language descriptions and advanced simulation software by constraining LLM outputs using a data schema (*i.e.*, format) and a modular component architecture, followed by deterministic validation to ensure correctness and reproducibility of the outputs.

**Results:** Using Monte Carlo eXtreme (MCX) – a widely used photon transport simulator – as an example, we showcase the capability of the proposed framework to convert user descriptions to structured simulation inputs. Benchmarked using 33 diverse natural language simulation descriptions, our LLM interface, MCX-LLM, achieves 98% accuracy and 99% repeatability, with an average processing time of 8.96 seconds per prompt. The framework also successfully handles various linguistic styles and diverse simulation settings, achieving a 100% success rate on 20 unconstrained real-world prompts. With only minor adjustments, our LLM interface also produces valid inputs for a finite-element-based diffusion solver to demonstrate generality towards other optical simulators.

**Conclusions:** By combining LLMs’ capability for textual data comprehension with structured constraints, this work provides a pathway to making complex scientific tools accessible while ensuring the reliability and technical correctness required for rigorous scientific research. MCX-LLM has been integrated with MCX Cloud accessible at https://mcx.space/cloud.

## 1 Introduction

Physical simulations and the use of specialized data analysis software packages are common in the field of biophotonics research.^1–4^ To obtain physically accurate modeling results, users must have not only an in-depth understanding of the underlying physical principles, but also knowledge of software-specific settings and their associated underlying assumptions, implied approximations, and inherent limitations. As computational software becomes increasingly complex, designing and executing effective simulation or data analysis pipelines often requires users to dedicate significant time to learning the software tools through specialized training and familiarizing themselves with complex interfaces, parameter configurations, and domain-specific conventions.^5^ This creates a steep learning curve and barrier for novice users and slows exploratory workflows for computational modeling and simulation.^4^ Consequently, a widening accessibility gap emerges between the growing capabilities of increasingly powerful open-source tools and the ability of typical users to fully utilize them.^6^ Enabling high-level interactions that allow users to express scientific intent without requiring expert-level software expertise could substantially improve the accessibility and usability of scientific simulation software.

Many scientific software tools already employ different user interface designs to facilitate users of different experience levels.^5^ Command-line interfaces (CLIs) offer flexibility and are well suited for batch processing and automation, but require users to understand specific syntax and input parameters. Graphical user interfaces (GUIs) lower the barrier to entry but typically limit access to advanced functions and batch processing. Language-specific bindings – such as MATLAB or Python wrappers – provide greater control and extensibility,^2, 4^ but still require users to provide structured input via commands and have sufficient knowledge to specify them correctly. These constraints make it difficult for users to take full advantage of complex simulators or data processing packages, especially during exploratory stages of a project.

Optical forward and inverse modeling software provides a concrete example that highlights the challenges of interfacing with scientific simulation tools. Although many biophotonics simulation tools offer different user interfaces,^7–10^ users must specify a large array of physical and simulation parameters – including light source properties, detector configurations, medium optical properties, and input/output settings, among others. For simulations involving specialized hardware such as a graphics processing unit (GPU), determining appropriate computing configurations – such as the total number of threads and block sizes – can also have a major impact on the simulation efficiency.^9, 11^ Correctly configuring these parameters often requires careful understanding of the documentation and substantial domain knowledge. Unintentional mistakes in the input format, value, or range can lead to failed simulations or physically unrealistic results. As a result, users without significant experience may struggle to use the software effectively or to exploit its full computational capabilities.

Large language models (LLMs) are a special type of artificial neural network (ANN) that have demonstrated extraordinary capability of understanding sophisticated natural language communications in terms of text and graphics.^12, 13^ LLMs have experienced explosive growth in recent years due to advances in neural network architectures, model scaling, large-scale data availability, and hardware improvements.^14^ Modern LLMs, trained on massive datasets and scaled with billions of parameters, are able to perform tasks such as natural language based conversations, code generation, and image pattern recognition. However, LLMs also have important limitations that restrict their direct use in scientific applications. They can produce hallucinated or incorrect information,^12^ and their outputs are generally not deterministic.^15^ Because LLMs model probability distributions over token sequences rather than executing explicit symbolic or numerical algorithms, they often struggle with precise logical or numerical constraints.^16, 17^ These issues make it difficult to rely on LLMs alone for tasks that require strictly specified formats and precise numerical inputs, such as configuring scientific simulations.

LLMs have been integrated into a wide range of software applications. Examples include conversational agents for customer support,^18^ code generation assistants such as Claude (Anthropic, CA), and natural language interfaces for data analysis.^19^ These applications leverage LLMs’ ability to process natural language to streamline workflows and improve productivity. However, due to the aforementioned challenges, their application for interfacing scientific software remains quite limited.^20, 21^ Scientific software typically requires strict input formats and precise numerical parameters, while LLM outputs often lack guarantee of correctness and adherence to physical constraints. Unlike customer support or general-purpose code generation, where approximate or conversational responses are often sufficient, scientific simulations require configurations that are syntactically valid, semantically complete, physically meaningful, and reproducible. Even small structural errors can cause simulations to fail or produce misleading results.

In this paper, we present a generalizable framework aiming at bridging the gap between LLMs and quantitative scientific simulation software.^22^ Rather than relying on the LLM to produce a perfect output in a single pass, we separate the problem into two parts: the LLM handles the flexible interpretation of natural language inputs, while a deterministic code pipeline enforces the schema constraints required by the simulator. As a concrete implementation of this framework, we introduce MCX-LLM, an open-source tool that integrates LLMs with the optical forward simulation toolbox Monte Carlo eXtreme (MCX)^9^ to enable natural-language-based simulation setup. MCXLLM employs Pydantic^23^ – a Python library for type enforcement – to define a human-readable schema that constrains the LLM’s output structure, and LangChain^24^ to construct prompt templates tailored for MCX simulations. A deterministic post-processing pipeline then parses and validates the JavaScript Object Notation (JSON)-formatted outputs to ensure that the generated configurations are consistent, well-structured, and compliant with MCX’s input format. The framework can be easily adapted to support diverse optical simulators^25–27^ and image reconstruction software pipelines.^1, 2, 28^

In the following section, we describe in detail the prompt template design, its integration with LLMs, and the subsequent deterministic schema-enforced input file generation process. In the Results section, we evaluate MCX-LLM across five aspects: (1) performance in terms of speed, accuracy, and repeatability under prompts of varying complexity, (2) robustness to language variation, (3) effectiveness in simulated real-world usage scenarios with human-composed prompts, and (4) comparison with commercially available general-purpose LLM-based chatbots, and (5) comparison with supervised fine-tuning as an alternative model-adaptation strategy. Finally, we discuss the limitations of this approach and outline potential future directions for interfacing scientific simulations with natural language.

## 2 Methods

The overall workflow of MCX-LLM is illustrated in Fig. 1. Our approach consists of three main components: (1) schema-guided prompt engineering to create composed LLM inputs, (2) deterministic post-processing to ensure MCX compatibility, and (3) integration with MCX Cloud for accessible deployment. In this work, we use MCX as a representative case study of this framework and evaluate its effectiveness through five comprehensive tasks.

**Fig 1:**
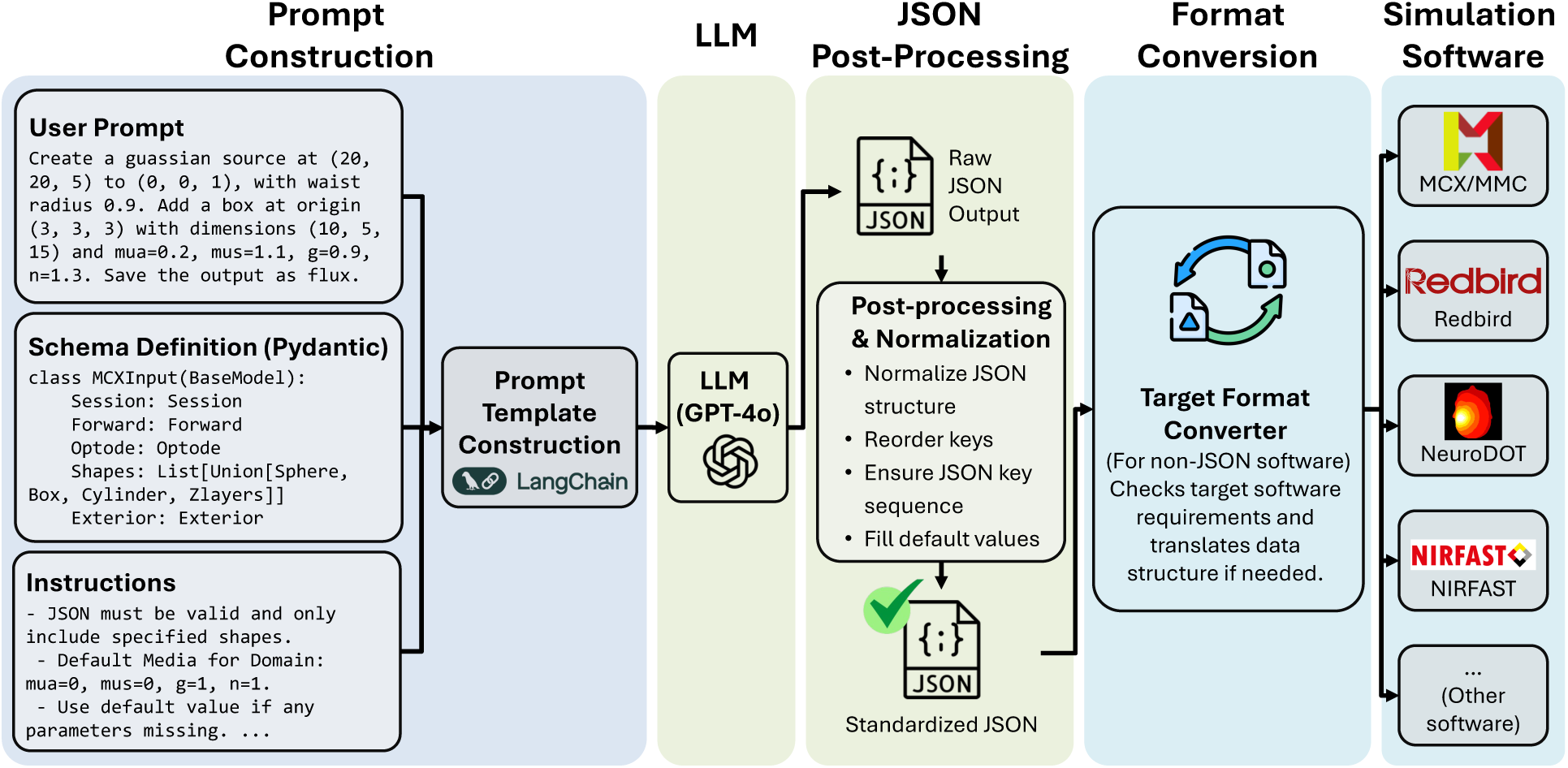
Overall workflow of MCX-LLM. A user’s natural language description of the simulation (sample snippet shown) is combined with a predefined JSON schema and detailed instructions (i.e., the system prompt) using a systematic prompt template (via LangChain). The combined prompt is passed to an LLM application programming interface (API) – in this case GPT-4o (OpenAI). The raw output from the LLM is subsequently processed by a deterministic post-processing script and optional converters to generate application-specific input files.

### 2.1 Schema-guided prompt engineering for MCX-LLM integration

To establish a connection between natural language prompts and the MCX simulator, we utilize JSON – a widely used structured data exchange format used by MCX^29^ – as the intermediate representation. Schema-guided prompt engineering techniques are used to convert natural language descriptions to structured input data usable by MCX.

Because LLMs are commonly implemented as autoregressive language models,^30^ they do not inherently guarantee strict structural consistency in their outputs. This often leads to unbalanced syntax, missing fields, or hallucinated content in the generated JSON data.^13^ To mitigate these issues, we design a schema to constrain the conversion process and configure the LLM parameters to produce outputs with reduced volatility. Specifically, we define a schema template based on MCX’s JSON input format using Pydantic,^23^ a Python library for type enforcement. The schema is supplied as one of the LLM inputs, serving as an explicit constraint that guides the model to produce syntactically valid and structurally consistent outputs.

Rather than reproducing the entire MCX schema – which would exceed the model’s token limit and include many unused parameters – we provide a subset schema focusing on the most commonly used fields. We also rename and reformat variables to be more self-explanatory and human-readable. This design balances schema completeness with input length, ensuring interpretability within the LLM’s context window^31^ while maintaining the essential structure needed for accurate simulation specification.

In addition to the schema, detailed textual instructions, known as the “system prompt”, are provided to guide the LLM’s responses. These instructions specify that the LLM must (1) produce valid JSON output, (2) preserve all required keys and assign default values when parameters are unspecified, and (3) correctly interpret source, detector, and domain properties, among other constraints. This strategy improves output reliability while maintaining flexibility to handle diverse user prompts across different phrasing styles and levels of technical detail.

The user’s natural language simulation description, JSON format template, and the system prompt are programmatically assembled using the PromptTemplate mechanism provided by LangChain.^24^ This process constitutes the core of our prompt engineering design, ensuring structured conditioning of the model’s generative behavior.

For this work, we ran all benchmarks using the GPT-4o model, accessible via the OpenAI API. The “temperature” parameter – a key setting common to most LLMs dictating the “creativity” of its generative responses – is set to 0 to greatly reduce stochastic variation and enforce deterministic outputs. This configuration maximizes reproducibility across repeated runs with identical prompts. We want to highlight that our prompting strategy is applicable to all LLM frameworks, including open-weight models.

After prompt assembly, the LLM generates a JSON-formatted response representing the requested simulation configuration. This output is then passed to a post-processing pipeline, described in the next subsection, which further aligns the generated data with MCX’s accepted input conventions.

### 2.2 Post-processing

To ensure that the LLM-generated JSON output is fully compatible with MCX, we implement a deterministic, code-based post-processing pipeline to verify and correct the model’s outputs. This pipeline consolidates the potential discrepancies between our simplified, human-readable schema and MCX’s native format requirements. It consists of several sequential steps that transform the LLM output into a fully MCX-compliant configuration.

First, we enforce structural consistency by renaming keys from our simplified schema to match MCX’s native conventions, standardizing key order according to MCX’s requirements, and reorganizing all media properties under the `Domain` input section. Next, we normalize parametrically described domain inclusions (spheres, cylinders, layered media, etc.) stored under the `Shapes` section, including reorganizing shape definitions and adding required shape parameters. The postprocessing step also inspects the LLM-generated string-valued specifiers, such as the output-type descriptor, and converts them to MCX-accepted values; it also fills in any missing fields using MCX’s default settings. Together, these operations produce a JSON configuration that is syntactically valid, semantically complete, and fully compatible with MCX.

This deterministic post-processing layer serves as a critical safeguard, ensuring that even when LLM output deviates slightly from the expected specification, the final JSON output remains robust and reproducible. This design illustrates how structured post-processing can bridge the gap between flexible natural language interfaces and the strict requirements of scientific simulation software.

### 2.3 Evaluation

We design five evaluation tasks to assess MCX-LLM’s performance as a natural language interface for quantitative Monte Carlo simulations. These tasks examine different aspects of the pipeline: general performance, handling of language variations, adaptability to diverse simulation tasks, performance relative to existing general-purpose LLMs, and comparison with supervised finetuning as an alternative model-adaptation strategy.

#### 2.3.1 Task 1 – accuracy, repeatability, and speed evaluation

To evaluate overall performance, we manually create 33 MCX simulation descriptions (i.e., prompts) in English, categorized into 8 groups based on complexity, with each group containing three to five descriptions. These groups evaluate performance for (1) single shape, (2) multi-shape, (3) output types and photon counts, (4) multi-shape and complex source, (5) time steps, (6) detectors, (7) optical property names, and (8) background domain. Specifically, the “single shape” category contains prompts featuring only one inclusion; “multi-shape” includes scenarios with multiple inclusions; “output types and photon counts” tests whether the model can handle various output format specifications (*e.g.*, flux, fluence, energy deposit, and momentum transfer) along with various simulation photon numbers; “multi-shape and complex source” includes prompts with multiple inclusions and complex source types that require setting additional parameters; “time steps” specify the starting time, ending time, or time-gate width of the simulation; “detectors” define the detector locations and sizes, which are optional in MCX simulations; “optical property names” describe the optical properties (*µ_a_*, *µ_s_*, *n*, *g*) by their full names (*i.e.*, absorption coefficient, scattering coefficient, refractive index, and anisotropy); “background domain” additionally defines the optical properties in the background to be different from the region outside the simulation domain.

Each prompt is executed 10 times, totaling 330 runs. We use three metrics: accuracy, repeatability, and speed. For accuracy, each run receives a score of 1 if the generated JSON output matches the expected result and 0 otherwise, with an average score computed per prompt. For repeatability, we first determine the most frequent output for each prompt across its 10 runs, then assign a score of 1 if a given run matches this most frequent output and 0 otherwise. Speed is measured as the average processing time per prompt from submission to the final output generation. Representative examples of simulated domains corresponding to different prompts are shown in Fig. 2.

**Fig 2:**
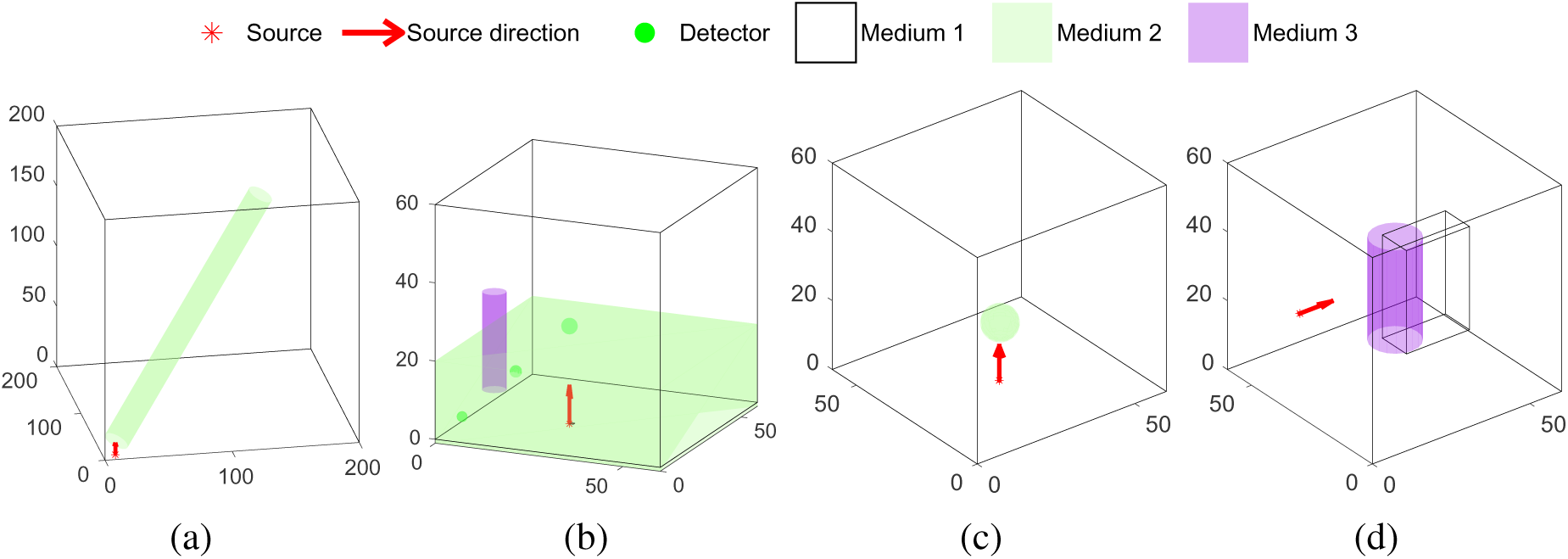
Representative MCX simulation domains generated by MCX-LLM from diverse simulation descriptions. Shown are examples with different geometries, embedded structures, detector placements, and source types, showing domains with (a) an isotropic source with a tilted cylindrical inclusion, (b) a two-layer structure with cylindrical inclusion and multiple detectors, (c) an isotropic source with a spherical inclusion, and (d) a hyperboloid source with a box and cylindrical inclusion. Colors are used to distinguish domain setting components (e.g., inclusions, detectors, and layered regions); un-shaded space is filled with medium label 1; red arrows indicate the source location and direction.

#### 2.3.2 Task 2 – language variation comprehension test

This task assesses how linguistic variation affects MCX-LLM’s output consistency. We specify a target configuration – a sphere within a two-layer structure (Fig. 3) – and provide a reference JSON file containing anticipated/ground-truth shape and simulation parameters. Four students with varying levels of familiarity with MCX are asked to independently write natural language simulation prompts describing this target configuration using their own phrasing and terminology. We then compare the JSON outputs generated by MCX-LLM from each student’s prompt to evaluate consistency and accuracy across different linguistic formulations of the same simulation intent.

**Fig 3:**
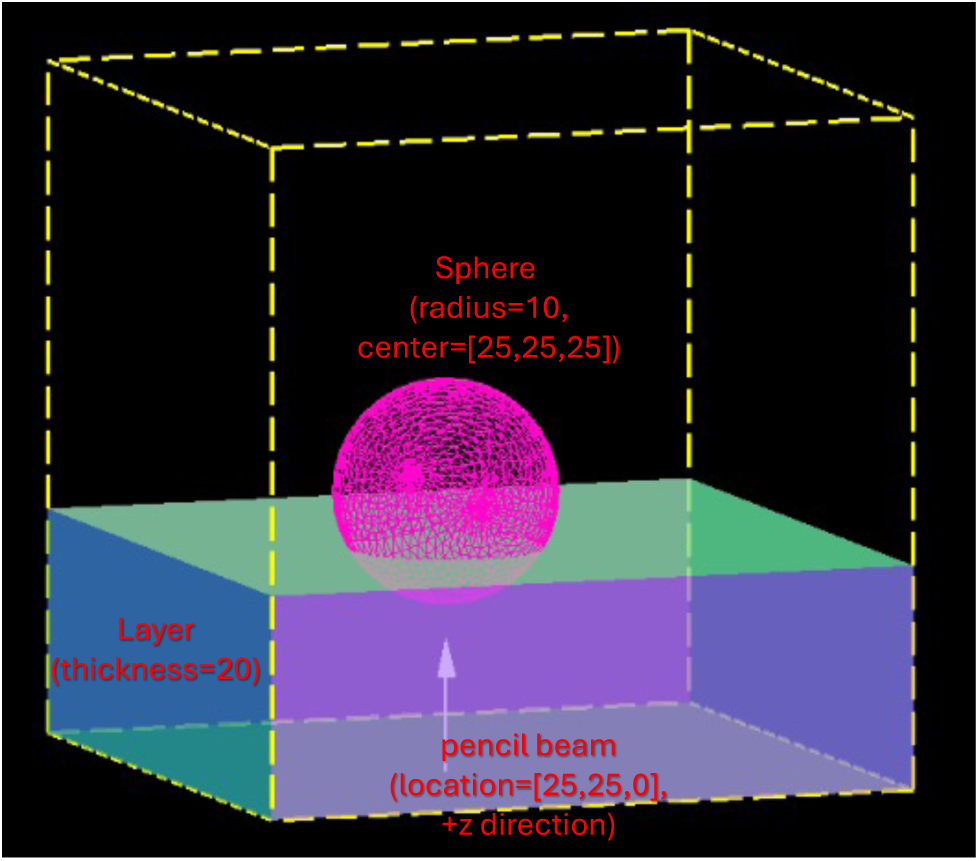
Rendering of the simulation domain used in the language variation comprehension test, illustrating a box domain containing a layered medium, a spherical inclusion, and a pencil-beam source.

#### 2.3.3 Task 3 – diverse simulation adaptability assessment

Because MCX-LLM is intended for use within the broad research community, it must have the ability to effectively handle diverse language preferences, technical backgrounds, and simulation configurations. To test the diversity of the simulations it can handle, each of four volunteers is asked to create five simulation prompts in English without restrictions on the type of simulations or their wording choices. This results in 20 prompts varying in language style, level of technical detail, and simulation complexity, offering a realistic basis for evaluating the pipeline’s ability to serve as a generalized interface across different communication styles and user expertise levels.

#### 2.3.4 Task 4 – baseline comparison with commercial LLM chatbots

To contextualize MCX-LLM’s performance, we compare it against three widely available commercial LLM-powered chatbots: ChatGPT,^32^ Claude,^33^ and Grok.^34^ Specifically, we test GPT-5-instant, Claude Sonnet 4.5, and Grok-4-fast. The same 33 prompts from Task 1 are used. To ensure a fair comparison, we provide each chatbot with the complete MCX JSON schema^29^ document, which strictly defines the type and meanings of every accepted input field, along with the same system prompt containing detailed instructions about the expected output structure. Each prompt is tested in an independent chat session to avoid memory effects between prompts and ensure consistent starting conditions.

#### 2.3.5 Task 5 – Supervised model fine-tuning

All commercial LLMs used in Task 4 are pretrained, and have no specific knowledge regarding the format and structure of MCX’s JSON input. To enhance the robustness of pretrained LLMs in generating MCX JSON inputs, we apply supervised fine-tuning – a technique that adapts a pretrained LLM using task-specific input-output examples so that the model can better follow instructions and generate outputs in the desired format^35, 36^.

For the fine-tuning experiment, each training sample consists of a natural-language MCX simulation description as the input and the corresponding ground-truth MCX-compatible JSON configuration from Task 1 as the target output. A 5-fold cross-validation is performed – the 33 manually curated prompts are divided into 5 non-overlapping groups, with 7 prompts each for 3 groups and 6 each for the remaining two groups; we use the prompts in each group in turn as the testing dataset and the remaining 26–27 prompts as the fine-tuning dataset, rotating across all 5 groups. The split is performed at the prompt level so that the test prompts are not seen during fine-tuning, allowing us to evaluate whether the fine-tuned model could be generalized to unseen MCX simulation descriptions. Because of the relatively small fine-tuning sample size, we record only the best result across the 5-fold testing as an indicator for any benefit due to additional fine-tuning.

### 2.4 Integration with MCX Cloud

To demonstrate the practical applicability of LLMs as generalized interfaces for scientific software, we integrate MCX-LLM into MCX Cloud,^29^ our existing web-based simulation platform. We add a new “LLM” tab where users can enter simulation prompts in plain language. These prompts are processed by MCX-LLM to generate MCX-compatible JSON configurations. After generation, users can review the resulting configuration in the “JSON” tab, visualize the simulation geometry in the “Preview” tab, or proceed directly to the “Run” tab to execute simulations using MCX Cloud’s existing infrastructure (Fig. 4). This integration demonstrates how natural language interfaces can be seamlessly incorporated into existing scientific software workflows without disrupting established functionalities or requiring significant platform redesign.

**Fig 4:**
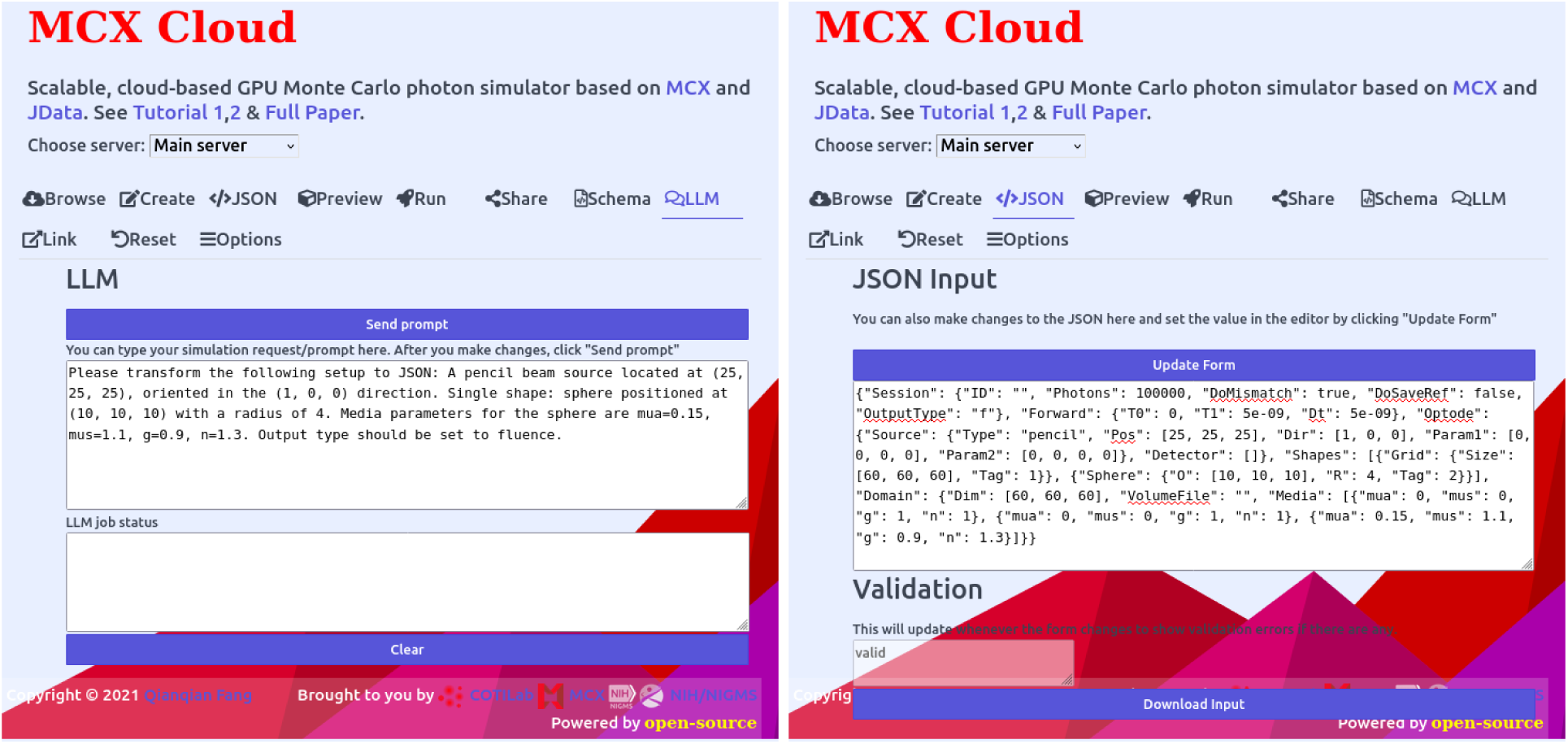
Integration of MCX-LLM into MCX Cloud. The left panel shows the “LLM” tab where users submit natural language simulation prompts, and the right panel shows the generated MCX-compatible JSON configuration in the “JSON” tab.

## 3 Results

### 3.1 Task 1 – accuracy, repeatability, and speed

Table 1 summarizes the performance metrics across all 8 prompt categories from 33 tested prompts. The overall accuracy is 0.98, repeatability is 0.99, and the average processing time is 8.96 seconds per prompt. Six categories – single shape, multi-shape, output types & photon counts, multi-shape & complex source, time steps, and detectors – achieve perfect scores (1.00) for both accuracy and repeatability. The “optical property names” category achieves an accuracy of 0.97 ± 0.06 and repeatability of 0.97 ± 0.06. The “define background domain” category shows the lowest performance, with an accuracy of 0.77 ± 0.40 and repeatability of 0.90 ± 0.17.

**Table 1:**
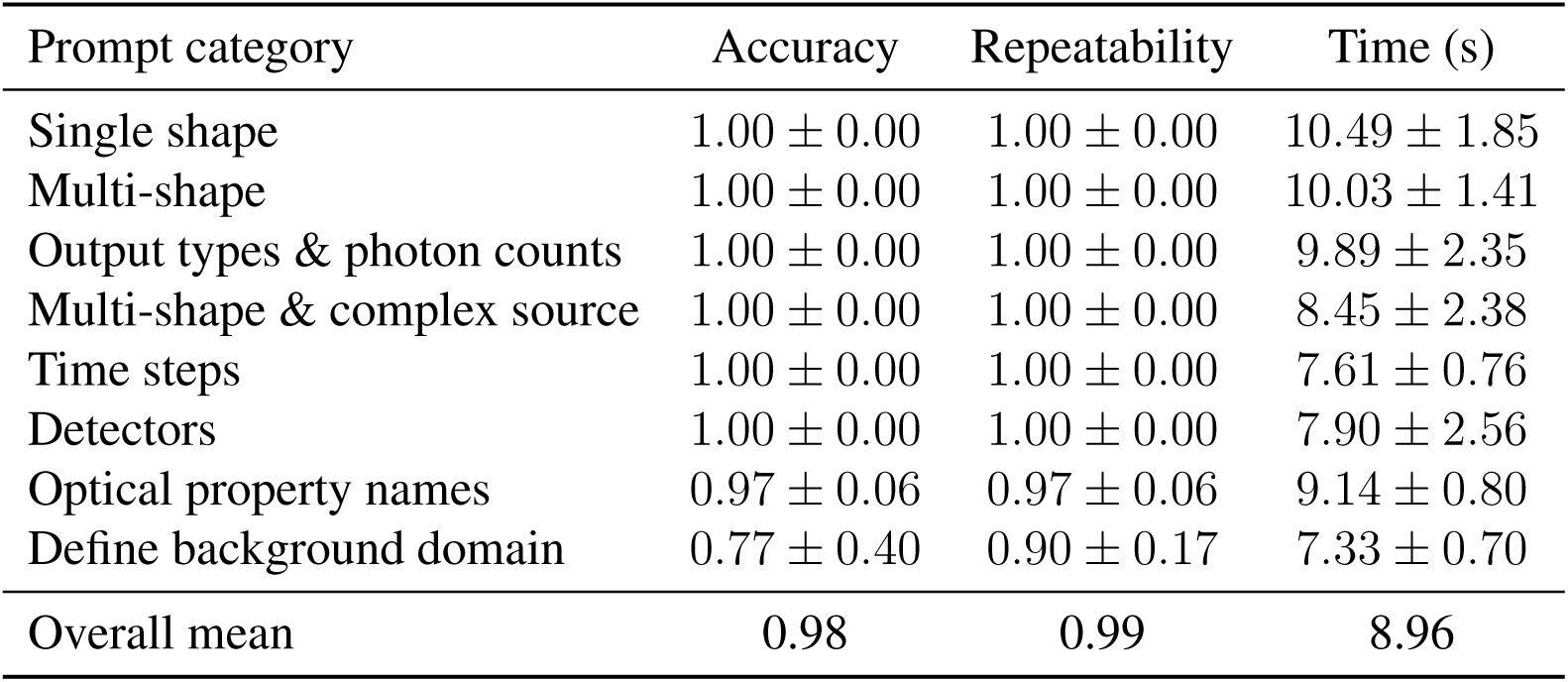
Performance metrics for MCX-LLM across 8 prompt categories. Values are reported as mean ± standard deviation over 10 repeated runs per prompt. Accuracy and repeatability are unitless scores between 0 and 1; time is measured in seconds.

### 3.2 Task 2 – language variation comprehension

Four volunteers (adult graduate students) independently generated natural language prompts describing identical simulation parameters. Table 2 shows excerpts from each subject’s prompt, along with word count and processing time. Prompt lengths range from 112 to 173 words, and processing times range from 7.90 to 11.30 seconds. The complete prompts used in this task are provided in Table S2 in the Supplementary Material. Despite substantial differences in phrasing and sentence structure, all four prompts produce identical JSON outputs.

**Table 2:** Excerpts from four subjects’ natural language prompts describing the same simulation task.

| Subject | Prompt preview | Words | Time (s) |
| --- | --- | --- | --- |
| S1 | “In this example, the MCX simulation domain consists of a spherical inclusion with a radius of 10 is centered at [25,25,25]...” | 133 | 11.30 |
| S2 | “Please prepare an MCX JSON file with the following settings: The whole domain is a 60 mm × 60 mm × 60 mm...” | 173 | 9.77 |
| S3 | “Can you generate a forward simulation to output the fluence? The domain is a box with x,y,z dimensions (60, 60, 60)...” | 130 | 9.26 |
| S4 | “Create an MCX simulation over a 60 by 60 by 60 voxelated domain, containing a spherical inclusion centered at [25, 25, 25]...” | 112 | 7.90 |

### 3.3 Task 3 – diverse simulation adaptability assessment

Table 3 summarizes the distribution of features across 20 English descriptions of diverse simulation settings from 4 volunteers. The complete prompts are provided in Table S3 in the Supplementary Material. MCX-LLM generates valid, executable JSON configurations for all 20 prompts, achieving a 100% success rate.

**Table 3:** Distribution of simulation task complexity across 20 unconstrained user prompts. The numbers in the “Range” column indicate the lower and upper bound of the respective feature in all tested prompts.

| Feature | Range | Coverage |
| --- | --- | --- |
| Geometric objects | 1–3 | Sphere, cylinder, box, layered structures |
| Source types | N/A | 6 types (pencil, Gaussian, cone, isotropic, disk, hyperboloid) |
| Detectors | 0–2 | 40% (8/20 prompts) include detectors |
| Layer structures | 0–3 | 40% (8/20 prompts) include layered media |
| Prompt length (words) | 48–201 | Mean: 108.40, std.: 48.30 |

### 3.4 Task 4 – comparison with commercial LLM-powered chatbots

We test three commercial LLM-powered chatbots – ChatGPT (GPT-5 Instant), Claude (Sonnet 4.5), and Grok (Grok-4 fast) – using the same 33 prompts from Task 1, resulting in 99 total responses (33 prompts × 3 chatbots). Only 6 responses (∼6%) produce MCX-acceptable JSON files, all of which are generated by Claude. ChatGPT and Grok each produce zero valid outputs.

Among the 93 invalid responses, the following error categories are identified (some responses contain multiple errors): missing default background (MCX requires the definition of the optical properties for label “0” – denoting the background medium) media properties (*n* = 81), incorrect output type specifications (*n* = 17), MCX schema documentation mixed into the output JSON (*n* = 8), overwriting between optical properties of different media (*n* = 4), and other structural or semantic errors (*n* = 8). These failure modes are categorized and compared against MCX-LLM in Table 4.

**Table 4:** Failure modes in LLM-generated MCX JSON configurations. Percentages represent failure rates for each tool; examples show errors in diff-style notation (red lines with “−” denote incorrect output; green lines with “+” denote the expected correct settings).

| Failure reason | MCX-LLM | Chatbots | Failure example (diff-style) |
| --- | --- | --- | --- |
| Incorrect default background properties | 2.4% (8/330) | 6.1% (6/99) | `"Media": [ - {"mua":0.001, "mus":1, "g":0.02, "n":1.38}] + {"mua":0, "mus":0, "g":1, "n":1},` |
| Violation of JSON schema | 0% (0/330) | 89.9% (89/99) | `"Media": [ + {"mua":0, "mus":0, "g":1, "n":1}, {"mua":0, "mus":0, "g":1, "n":1}, {"mua":0.2, "mus":1.5, "g":0.85, "n":1.45}]` |
| Contradicting user’s prompt | 0% (0/330) | 17.2% (17/99) | `"Photons": 100000, - "OutputType":"f",//Generated fluence ("f") + "OutputType":"x",//Expecting fluence-rate ("x")` |

### 3.5 Task 5 – Supervised fine-tuning

For consistency with Task 1, we evaluate the fine-tuned models using exact JSON-value matching against the held-out ground-truth configurations. The fine-tuning baseline shows uneven performance across the held-out test set. Across the 5-fold rotations of the testing datasets, the best fine-tuned GPT-4o-mini model produced 5 of the 7 matched prompts, compared to 0 matches when using the pre-trained GPT-4o-mini without fine-tuning. However, the fine-tuned GPT-4.1 and GPT-nano variants fail to yield any improvement compared to pre-trained models, suggesting strong sensitivity to the choice of base model and training stability under the limited-data setting.

### 3.6 Generalization towards other optical simulators

To evaluate the generalizability of the proposed framework beyond MCX, we apply it to convert user descriptions to the input data structure for Redbird^26, 27^ – a MATLAB toolbox with a finite-element method (FEM) based diffusion equation solver and a nonlinear diffuse optical tomograph (DOT) reconstruction pipeline. As shown in Fig. 5, given a natural-language prompt, MCX-LLM automatically generates a Redbird-compatible configuration and successfully produces the corresponding simulation output. For example, we prompt the system to generate a Redbird simulation with a box-shaped domain, a single homogeneous region, and a 5 × 5 grid of sources and detectors placed on opposite faces of the volume. With only minor modifications to the Pydantic schema and post-processing code – requiring fewer than 10 lines of code changes – we are able to produce a valid JSON file that converts directly into a Redbird configuration structure.

**Fig 5:**
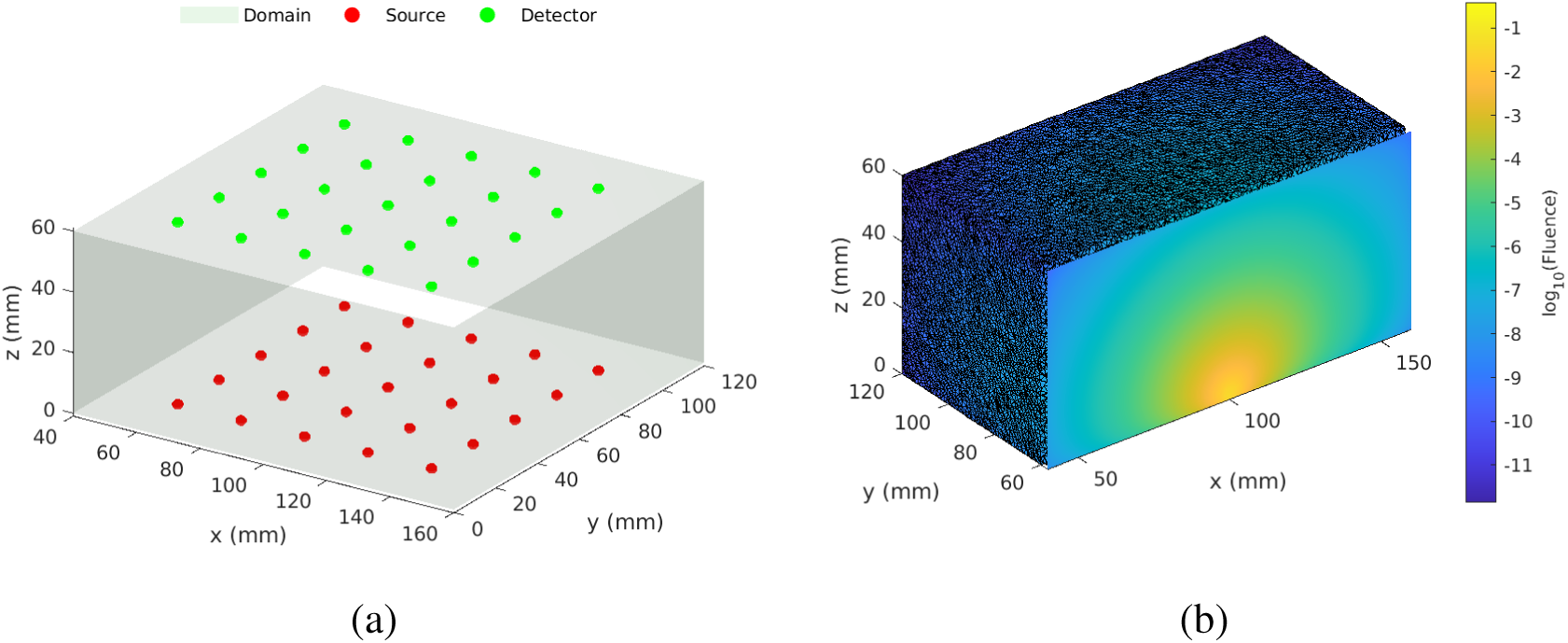
Extension of the proposed framework to Redbird: (a) the generated simulation setup from a natural-language prompt; (b) the corresponding simulation output produced by Redbird.

### 3.7 Evaluation with multilingual simulation descriptions

We further evaluate the use of non-English natural language input. Specifically, a subset (#1, #2, and #17) of the prompts used in Task 1 were translated into Chinese and re-processed using MCXLLM. Without changing any LLM settings, the resulting JSON outputs exactly replicate those generated from the respective English prompts. This result underscores a new benefit of using LLMs as scientific software interfaces – their inherent ability to comprehend and process multiple languages could significantly improve accessibility for researchers across different countries.

## 4 Discussion

In this work, we present MCX-LLM – a framework that bridges the gap between natural language based descriptions and advanced scientific simulations. Using MCX-LLM as the interface, we can accommodate users of diverse experience levels and linguistic preferences to create simulation configuration files that are not only valid in format, but also semantically correct and reproducible across language variations. Using our widely adopted GPU-accelerated Monte Carlo photon simulator, MCX, and our DOT forward and reconstruction platform, Redbird, we are able to create valid simulation configurations from several dozen tested user prompts spanning a wide range of simulation features. Below, we discuss the key findings from each evaluation task, followed by discussions towards broader implications and limitations of this approach.

The Task 1 results shown in Table 1 demonstrate the high accuracy (98%) and high repeatability (99%) of MCX-LLM across 330 runs. Six of the eight tested prompt categories obtain perfect scores for both metrics, indicating that the combination of schema-guided prompting and deterministic post-processing can reliably handle a wide range of simulation configurations. The “optical property names” and “define background domain” categories report slightly lower performance, suggesting that prompts requiring the LLM to map verbose natural-language terms (*e.g.*, full names of optical properties) to concise parameter keys, or to distinguish between background and domain-level media settings, remain unreliable. Nevertheless, even in the lowest-performing category, the repeatability at 90% indicates that most outputs are consistent across repeated runs. Because LLMs are known for producing nondeterministic outputs,^15^ these results suggest that our combination of Pydantic schema constraints and deterministic post-processing largely mitigates this variability for scientific computing applications.

The language variation comprehension test (Task 2) further confirms the robustness of the proposed framework. Despite substantial differences in phrasing and sentence structure across the four subjects (prompt length ranging from 112 to 173 words), MCX-LLM produces identical JSON outputs for all subjects. This consistency is critical for scientific reproducibility – different researchers describing the same experiment are expected to obtain the same simulation configuration regardless of their individual writing styles.

The diverse simulation adaptability assessment (Task 3) demonstrates that MCX-LLM can accommodate a wide range of simulation features, including both basic and advanced settings. All 20 unconstrained prompts, covering diverse geometric objects, source types, and detector configurations (Table 3), produce valid and readily usable JSON input files for MCX. This result is encouraging for the broad deployment of this tool to address real-world user needs.

The comparison with commercial LLM-based chatbots (Task 4) highlights the necessity of our schema-guided design. When provided with the same MCX schema and instructions, only 6% of responses from ChatGPT, Claude, and Grok produce valid MCX configurations, in contrast to MCX-LLM’s 98% accuracy. As shown in Table 4, the dominant failure mode among chatbots is generating JSON that violates the schema (89.9%), followed by not following the user’s prompt (17.2%) and not using default values (6.1%). These results underscore that general-purpose LLMs, even when supplied with detailed schema documentation, lack the structural constraints and validation mechanisms needed for reliable scientific software integration. In other words, simply providing schema documentation to a general-purpose LLM is insufficient to guarantee valid simulation configurations. The proposed schema-guided approach directly addresses this gap by constraining the LLM’s output space and applying deterministic corrections in post-processing.

The fine-tuning results in Task 5 further highlight the challenge of relying on model adaptation alone for scientific simulation input generation. Although the GPT-4o-mini fine-tuned model achieved some improvement over its pre-trained version, the overall performance remained highly dependent on the choice of base model and the coverage of the training examples. This behavior is generally consistent with the role of supervised fine-tuning, which can adapt model behavior to a target task but depends on representative training data.^35, 36^ MCX simulation configurations involve combinations of source types, geometric inclusions, optical properties, detector settings, time gates, output types, and default media definitions. A small training set is unlikely to cover this space sufficiently. As a result, fine-tuned models may still fail when an unseen prompt contains structures, terminology, or parameter combinations that are underrepresented in the training data. In contrast, the proposed schema-guided framework reduces the need for the LLM to internalize every simulator-specific rule. The LLM is used primarily to interpret the user’s intent, while deterministic post-processing enforces structural completeness, default values, and compatibility with the native MCX input format.

To make MCX-LLM accessible to broad user communities, we have integrated it with MCX Cloud.^29^ This integration allows users to design simulations by describing the setup in natural language, significantly lowering the barrier to entry for researchers unfamiliar with MCX’s configuration syntax. Moreover, the extension to Redbird^26, 27^ demonstrates that the proposed framework is not limited to MCX. Because the schema definition and post-processing rules are modular components, adapting the framework to a new simulation tool requires only defining the target schema and the corresponding mapping rules. In practice, modifying fewer than 10 lines of the Pydantic schema and corresponding post-processing code is sufficient to produce valid Redbird configurations from natural-language descriptions. This suggests a generalizable approach for connecting human language with machine-readable input formats required by a range of scientific simulation tools.

Contemporary prompt engineering strategies such as few-shot prompting^30, 37, 38^ represent new avenues for improving LLM-generated simulation inputs without modifying model weights. In few-shot prompting, a small number of input-output demonstrations are included in the prompt to guide the model through in-context learning.^30^ In our exploratory tests, we observed a substantially improved success rate when providing detailed descriptive instructions (*>*1,000 words) and representative examples (i.e., shots) composed of description-JSON pairs, confirming the strong potential of LLMs for natural-language comprehension and structured text generation. However, relying on prompt-only approaches inherently risks producing unreliable output for error-sensitive applications such as scientific simulation. Their performance can vary across model families, model versions, decoding settings, and prompt wording choices. Incomplete descriptions, ambiguous terminology, or simulation requests that differ from the examples included in the prompt may further hamper the reliability of the result. Moreover, an LLM may have difficulty strictly following long schemas and complex instructions, leading to missing default media properties, incorrect output-type mappings, media-ordering errors, or hallucinated fields. While such errors may be tolerable or manually correctable in conversational applications, they can cause failed simulations or incorrect results in MCX. Therefore, prompt engineering is valuable for improving LLM behavior, but schema constraints and deterministic validation remain necessary for achieving the correctness and reproducibility required by scientific simulation software.

Several limitations should be noted. First, the current version of MCX-LLM does not support all features available in MCX. Because simulation domains are specified using natural language, the current implementation restricts domains to a predefined set of geometric shapes (under the “Shapes” JSON input section), which limits extensibility to more complex or patient-specific geometries. To support voxel-based volumes in future work, two approaches may be considered: (1) dynamically querying and downloading pre-made simulation volumes from NeuroJSON.io^39^ – a large repository of freely accessible medical images, segmentation volumes, and mesh models – and (2) referencing the user’s uploaded custom volume data (already supported in MCX Cloud) in the simulation.

## 5 Conclusion

In summary, we present MCX-LLM – a framework that bridges the gap between natural language and scientific simulation software by combining LLM’s flexibility with schema-guided constraints and deterministic post-processing. Using MCX as a representative case study, we demonstrate that this approach achieves 98% accuracy and 99% repeatability across diverse simulation prompts while maintaining an average processing time under 9 seconds. The framework successfully handles language variation, unconstrained user inputs, and multilingual prompts, and substantially outperforms general-purpose chatbots in generating valid simulation configurations. We further demonstrate the generalizability of this approach through its extension to Redbird with minimal code modifications.

## Code, data, and materials availability

MCX-LLM has been integrated with MCX Cloud at https://mcx.space/cloud and is freely available to global biophotonics researchers.

## Acknowledgments

This research is supported by the National Institutes of Health (NIH) National Institute of General Medical Sciences (NIGMS) grant R01-GM114365, National Institute of Biomedical Imaging and Bioengineering grant R01-EB026998, and the National Institute of Neurological Disorders and Stroke (NINDS) grant U24-NS124027. This study is specifically aimed at investigating LLMs for converting user descriptions into structured scientific simulation settings. Various LLM models have been tested with results summarized in the Results section. Additionally, the authors used Claude (Anthropic, CA) and ChatGPT (OpenAI, CA) for language editing, grammar correction, and clarity improvement during manuscript preparation.

