## Supplementary Material for "Using large language models for enhancing accessibility for Monte Carlo photon transport simulations and beyond"

#### S1. Complete Prompts Used for Task 1

Table S1: Complete natural language prompts used for the Task 1 accuracy, repeatability, and speed evaluation.

| ID | Prompt |
|---|---|
| | **Single shape** |
| Prompt 1 | Please transform the following setup to JSON: A pencil beam source located at (25, 25, 25), oriented in the (1, 0, 0) direction. Single shape: sphere positioned at (10, 10, 10) with a radius of 4. Media parameters for the sphere are mua=0.15, mus=1.1, g=0.9, n=1.3. Output type should be set to fluence. |
| Prompt 2 | Generate JSON format for an isotropic source located at (15, 15, 0), direction (0, 0, 1). Add one cylinder stretching from (5, 5, 5) to (5, 5, 30) with a radius of 2. Cylinder media values are mua=0.2, mus=1.5, g=0.85, and n=1.45. Set output type to energy deposit. |
| Prompt 3 | Convert this configuration into JSON: Place an isotropic source at (10, 10, 0), pointing to (0, 1, 0). Include a box originating at (2, 2, 2) with dimensions (8, 8, 8), with media parameters mua=0.1, mus=1.0, g=0.8, and n=1.3. The output type should be output flux. |
| Prompt 4 | Format as JSON: A Gaussian source situated at (30, 30, 0), pointed along (0, 0, 1), with a waist radius of 3. Add Z-layers at depths from 0 to 10. Layer media properties are mua=0.2, mus=1.2, g=0.75, n=1.5. |
| Prompt 5 | Request JSON formatting: Use a cone source located at (20, 20, 0) with a half-angle of 0.3 radians. Single sphere positioned at (10, 10, 10) with radius 6. Sphere media properties include mua=0.25, mus=0.9, g=0.85, and n=1.4. Set output type to fluence. |
| | **Multi-shape** |
| Prompt 6 | Do a simuluation with hyperboloid source at (15, 15, 0) with a waist radius of 2.5, focus distance of 4, and Rayleigh range of 12. Include a layer at 0 with a thickness of 15 along z axis, and add a sphere at (10, 10, 10) with radius 5. Sphere media: mua=0.1, mus=0.9, g=0.85, n=1.3. Layer media: mua=0.2, mus=0.9, g=0.95, n=1.33. Set output type to energy deposit. |
| Prompt 7 | Create a guassian source at (20, 20, 5) to (0, 0, 1), with waist radius 0.9. Add a box at origin (3, 3, 3) with dimensions (10, 5, 15) and mua=0.2, mus=1.1, g=0.9, n=1.3. Save the output as flux. |
| Prompt 8 | Convert to JSON: Cone source at (5, 5, 0) with half-angle of 0.25 radians, pointed along (1, 0, 0), shapes include a cylinder from (10, 10, 10) to (10, 10, 25) radius 3, and Z-layers at 0 with depth 20. Media: Cylinder mua=0.1, mus=0.6, g=0.85, n=1.4; Zlayer mua=0.4, mus=0.83, g=0.9, n=1.3. Use energy deposit. |
| Prompt 9 | Structure as JSON: Use an isotropic source at (30, 30, 0) to (0, 1, 0) with two shapes: a box at (5, 5, 5) size (10, 5, 15), and a sphere centered at (15, 15, 15) radius 6. Media: Box has mua=0.18, mus=0.75, g=0.8, n=1.3; sphere has mua=0.22, mus=0.9, g=0.9, n=1.4. Set output type to fluence. |

| | |
|---|---|
| **Output types & photon counts** | |
| Prompt 10 | Structure as JSON: Planar source positioned at (10, 10, 10). The planar surface is defined by two edge vectors: one extending by (5, 5, 5) from the source position, and the other extending by (1, 1, 1), oriented at (0, -1, 0). Single shape: a sphere at (20, 20, 20) with radius 8 and media properties mua=0.15, mus=0.7, g=0.85, n=1.37. Set output type energy deposit and photon number 400000. |
| Prompt 11 | Convert the following into JSON: Isotropic source at (5, 5, 5) to (1, 1, 1). Shapes include a box from (8, 8, 8) with size (5, 5, 20), and a layer is with thickness 15. Box is mua=0.1, mus=0.5, g=0.8, n=1.5, and layer is mua=0.15, mus=1.1, g=0.9, n=1.3. Choose output type flux. The simulation is with 3000000 photons. |
| Prompt 12 | Hyperboloid source at (10, 10, 0) to (0, 1, 0) with a waist radius of 1.5, focus distance 2, and Rayleigh range 10. Shapes include a box at (15, 15, 15) with size (8, 8, 8) and media mua=0.2, mus=0.7, g=0.85, n=1.45. Set photon counts to 1e7 and output energy. |
| Prompt 13 | Consider 500000 photons of a pencil beam source at 20, 20, 0, pointing at 1, 2, 0. With Two shapes in the domain: a cylinder from (10, 10, 0) to (10, 10, 30) with radius 3, the optical propety is mua=0.18, mus=0.85, g=0.88, n=1.4. And a sphere at (20, 20, 20) with radius 6 with mua=0.1, mus=0.7, g=0.9, n=1.3. Set output type flux. |
| Prompt 14 | Format as JSON: Cone source at (30, 15, 15) with a half-angle of 0.2 radians, oriented to (-1, 0, 0). Include a box at (8, 8, 8) with dimensions (6, 6, 6) and Z-layers range between 0 and 25. Box is mua=0.2, mus=0.75, g=0.88, n=1.36 and z-layer is mua=0.1, mus=0.8, g=0.9, n=1.33. Set output type fluence with photon count 600000. |
| **Multi-shape & complex source** | |
| Prompt 15 | Convert to JSON: Use a planar source at (15, 15, 0), oriented along (0, 1, 0). The plane is spanned by two vectors starting at the source point: (0, 1, 0) and (1, 0, 0). Define three shapes: a sphere at (10, 10, 10) with radius 5, a cylinder extending from (20, 20, 5) to (20, 20, 25) with radius 2, and a box at (5, 5, 5) sized (8, 8, 8). Sphere media is mua=0.2, mus=0.7, g=0.85, n=1.38; cylinder media is mua=0.1, mus=1.0, g=0.9, n=1.4; box media is mua=0.15, mus=0.6, g=0.87, n=1.5. Choose output type fluence. |
| Prompt 16 | Generate JSON structure: Gaussian source at (15, 15, 15), directed toward (0, 1, 1) with a waist radius of 1.5. Include two shapes: a sphere at (25, 25, 25) with radius 5, and a box at (10, 10, 10) with size (5, 5, 10). Sphere media properties: mua=0.2, mus=0.85, g=0.9, n=1.33; box media: mua=0.1, mus=1.1, g=0.88, n=1.37. Choose output type flux. |
| Prompt 17 | Cone source positioned at (10, 10, 0) with a half-angle of 0.25 radians. Add a Z-layer starting from, 0 to 20, along with a cylinder starting at (5, 5, 5) and ending at (5, 5, 25), radius 2. Cylinder media is mua=0.2, mus=0.6, g=0.85, n=1.45 and z-layer media is mua=0.1, mus=1,2, g=0.95, n=1.2. |
| Prompt 18 | Use a hyperboloid source at (20, 10, 10), pointing at (1, 0, 1), focus distance 3, rayleigh range 5, and waist radius 2.3. Define a sphere at (10, 10, 10) with radius 6, a box at (15, 15, 15) with dimensions (5, 10, 5), and Z-layers from 0 to 10. Sphere media: mua=0.12, mus=0.8, g=0.87, n=1.35; box media: mua=0.2, mus=0.9, g=0.9, n=1.4, and z-layer media: mua=0.1, mus=0.85, g=0.91, n=1.22 |
| Prompt 19 | Generate a isotropic source located at (50, 50, 0), pointing at (0, 0, 1) in a $100^3$ cube domain. A sphere is at (30, 30, 30) with radius 15 (mua=0.18, mus=0.85, g=0.88, n=1.4), and another sphere at same location but with radius 20 (mua=0.2, mus=0.75, g=0.88, n=1.36). Add two z layers, one is spanning from 0 to 20 (mua=0.1, mus=1.0, g=0.9, n=1.4), and the other one is from 21 to 40 (mua=0.15, mus=1.1, g=0.9, n=1.3). |
| **Time steps** | |

| | |
|---|---|
| Prompt 20 | Convert this configuration to JSON: Pencil source at (10, 10, 10) directed toward (1, 0, 0). Shapes include a sphere at (15, 15, 15) with radius 4, media mua=0.15, mus=0.9, g=0.85, n=1.35. Set Dt to 1e-09 and output type to energy deposit. |
| Prompt 21 | Generate JSON format: Gaussian source at (20, 20, 0) with waist radius 2.5 and directed to (0, 1, 0). Include a box at (5, 5, 5) with dimensions (10, 5, 5) and media mua=0.12, mus=1.2, g=0.88, n=1.34. Set Dt to 2e-09 and output type fluence. |
| Prompt 22 | Format as JSON: Gaussian source located at (30, 0, 10), directed along (1, 1, 0) with waist 2.5. Define two shapes: a box from (3, 3, 3) with size (10, 5, 15), and a sphere at (20, 20, 20) radius 4. Box media is mua=0.25, mus=1.2, g=0.9, n=1.36; sphere media is mua=0.1, mus=0.9, g=0.85, n=1.33. Set Dt to 2.5e-09. |
| Prompt 23 | Perform a simulation of isotropic source at (10, 10, 0), pointing towards (0, 0, 1) in a $200^3$ cube domain. The inclusion object is a cylinder from (10, 10, 10) to (150, 150, 150), radius 10, with optical properties: mua=0.2, mus=1.1, g=0.85, n=1.45. Set the Dt to 5e-8, and photon counts to 1e8. |
| **Detectors** | |
| Prompt 24 | Create JSON: Gaussian source at (15, 15, 0) with waist radius 2.5. Add a detector at (10, 10, 10) with radius 1.5. Define a sphere at (20, 20, 20) with radius 6 and media mua=0.1, mus=0.9, g=0.85, n=1.33. Set Dt to 3e-09 and output type fluence. |
| Prompt 25 | Convert to JSON: Cone source at (5, 5, 5) to (1, 0, 0) with a half-angle of 0.4 radians. Detectors are placed at (10, 10, 0) with radius 2 and (15, 15, 15) with radius 1. Add a cylinder from (10, 10, 10) to (10, 10, 30) with radius 3, and media mua=0.2, mus=0.8, g=0.87, n=1.4. Set Dt to 6e-09 and photon count 400000. |
| Prompt 26 | Consider a pencil beam at (10, 0, 10), pointing at (0, 1, 0) , with detectors at (8, 8, 8) radius 1.5 and (12, 12, 12) radius 0.9. Shapes include a box at (5, 5, 5) size (8, 8, 8), media mua=0.3, mus=1.0, g=0.85, n=1.45, and a cylinder from (15, 15, 5) to (15, 15, 25) radius 2, media mua=0.2. mus=1.3, g=0.9, n=1.23. |
| Prompt 27 | Create JSON structure: Planar source positioned at (25, 25, 0), oriented (0, 0, 1). Let the plane be defined by two edges coming out from that point: one in the (1, 0, 0) direction and the other in the (0, 1, 0) direction. Detectors include (5, 5, 5) with radius 1.3, (15, 15, 15) with radius 1.5, and (25, 25, 25) with radius 2. Add a Z-layer setup at depths 0 with a thickness of 20, and a cylinder from (10, 10, 10) to (10, 10, 35) with radius 3. Cylinder is mua=0.2, mus=0.85, g=0.88, n=1.37, and zlayer is mua=0.14, mus=0.78, g=0.94, n=1.2 |
| **Optical property names** | |
| Prompt 28 | Please transform this setup into JSON: Set a Gaussian source positioned at (10, 10, 10) with a waist radius of 1.5, pointing toward (1, 0, 0). Include a sphere centered at (20, 20, 20) with a radius of 4, and assign it the following optical properties: an absorption coefficient of 0.2, scattering coefficient of 1.0, anisotropy of 0.85, and a refractive index of 1.34. Place a detector nearby at (10, 10, 10) with a radius of 1.0. |
| Prompt 29 | Generate JSON: Use an Isotropic source at (5, 5, 5). Add a cylinder extending from (10, 10, 5) to (10, 10, 25) with a radius of 3 and a box at (8, 8, 8) with dimensions (6, 6, 6). Set the cylinder's optical properties to an absorption coefficient of 0.15, a scattering coefficient of 0.8, an anisotropy of 0.8, and a refractive index of 1.37. For the box, use an absorption coefficient of 0.05, scattering coefficient of 0.9, anisotropy of 0.85, and refractive index of 1.33. |
| Prompt 30 | Generate JSON: Use an Isotropic source at (5, 5, 5). Add a cylinder extending from (10, 10, 5) to (10, 10, 25) with a radius of 3 and a box at (8, 8, 8) with dimensions (6, 6, 6). Set the cylinder's optical properties to an absorption coefficient of 0.15, a scattering coefficient of 0.8, an anisotropy of 0.8, and a refractive index of 1.37. For the box, use an absorption coefficient of 0.05, scattering coefficient of 0.9, anisotropy of 0.85, and refractive index of 1.33. |
| **Define background domain** | |

| | |
|---|---|
| Prompt 31 | Create a MCX simulation with background properties mua=0.005, mus=1, g=0.01, n=1.37. The isotropic source is at 30, 30, 0, pointing to 0, 0, 1. A sphere is centered at 30, 30, 15, with radius 5, media is mua=0.18, mus=0.85, g=0.88, n=1.4. |
| Prompt 32 | Generate a JSON structure: Hyperboloid source at (0, 30, 30) to (1, 0, 0), focus distance 2, rayleigh range 4, and waist radius 1.3. The background is mua=0.001, mus=1, g=0.02, n=1.38. Two shapes in the domain, a box at (30, 25, 10), size (20, 10, 30), with mua=0.25, mus=1.2, g=0.85, n=1.3; and a cylinder two ends at (30, 30, 10) and (30, 30, 40), radius 7, with mua=0.2, mus=0.5, g=0.8, n=1.25. |
| Prompt 33 | Do a simulation with background mua=0.2, mus=10, g=0.9, n=1.37 and size 100*100*100. A pencil beam source located at (50,50,0), directed to (0,0,1). Inclusion is a zlayer from 0 to 10, with properties mua=0.02, mus=10, g=0.9, n=6.85. |

### S2. Complete Prompts Used for Task 2

Table S2: Complete natural language prompts from four subjects used for the Task 2 language variation comprehension test.

| Subject ID | Prompt |
|---|---|
| S1 | In this example, the MCX simulation domain consists of a spherical inclusion with a radius of 10 centered at [25,25,25]. Additionally, a layer spans from [0,0,0] to [60,60,20].<br>The optical properties for the first layer, second layer, and the inclusion are as follows:<br>• Inclusion: $\mu_a = 0.15$, $\mu_s = 1.1$, $g = 0.9$, and $n = 1.3$.<br>• The layer: $\mu_a = 0.01$, $\mu_s = 1.2$, $g = 0.75$, and $n = 1.5$.<br>The light source is a pencil beam positioned at [25,25,0], emitting light in the $+z$ direction. A total of $10^5$ photons are launched to compute the light fluence. The maximum time of flight simulated for each photon is 5 ns, with a single time gate used for the simulation. |
| S2 | Please prepare an MCX JSON file with the following settings: The whole domain is a 60 mm × 60 mm × 60 mm volume with a 1 mm edge length voxel size. Place a spherical object of radius 10 mm at position [25,25,25] mm with optical properties of $\mu_a = 0.15$ mm$^{-1}$, $\mu_s = 1.1$ mm$^{-1}$, $g = 0.9$, and $n = 1.3$. Set the rest of the domain to have optical properties of $\mu_a = 0$, $\mu_s = 0$, $g = 1$, and $n = 1$. From a z-position of 0 mm to 20 mm, set the optical properties of the domain to have $\mu_a = 0.01$ mm$^{-1}$, $\mu_s = 1.2$ mm$^{-1}$, $g = 0.75$, and refractive index $n = 1.5$.<br>Run the simulation with 100,000 photons. The source is a point source centered at [25,25,0] mm propagating in the direction of [0,0,1] and simulated over a total simulation time of $5 \times 10^{-9}$ seconds. Provide the output as volumetric fluence. |
| S3 | Can you generate a forward simulation to output the fluence? The domain is a box with x,y,z dimensions (60, 60, 60). The coefficients mua and mus are 0 and g and n are 1. Inside the box is a sphere centered at 25,25,25 with a radius of 10. Please set the sphere coefficients, mua to be 0.15, mus to be 1.1, g is 0.9 and n is 1.3. Please also set the z layers 0 through 20 of the box to have the following coefficients, mua is 0.01, mus is 1.2, g is 0.75, n is 1.5. The source is a pencil beam with direction (0,0,1) at position (25,25,0), launching 100000 photons. Thank you |

| Subject ID | Prompt |
|---|---|
| S4 | Create an MCX simulation over a 60 by 60 by 60 voxelated domain, containing a spherical inclusion centered at [25, 25, 25], with a radius of 10, and absorption coefficient 0.15/mm, scattering coefficient 1.1/mm, anisotropy 0.9 and refractive index 1.3; the domain also contains a z-layer object between z=0 and 20, filled with absorption coefficient 0.01/mm, scattering coefficient 1.2/mm, anisotropy 0.75 and refractive index 1.5. A pencil beam source is located at [25, 25, 0] pointing in the direction of [0, 0, 1]. Please run 1e5 photon packets for a maximum time window of 5e-9 seconds, and output fluence. |

### S3. Complete Prompts Used for Task 3

Table S3: Complete unconstrained natural language prompts used for the Task 3 adaptability assessment.

| Subject ID | Prompt ID | Prompt |
|---|---|---|
| S1 | Prompt 1 | I would like to simulate an isotropic light source within a spherical inclusion. The inclusion is centered at [30,30,30] with a radius of 30. The light source is positioned at the center of the sphere. The optical properties of the sphere are defined as $\mu_a = 0.05$, $\mu_s = 10$, $g = 0.9$, and $n = 1.3$. |
| | Prompt 2 | I would like to simulate the traversal of a light beam through a spherical lens. The lens is modeled as a spherical inclusion with optical properties $\mu_a = 0.0$, $\mu_s = 0.0$, $g = 1.0$, and $n = 1.4$. It is centered at [30 30 30] with a radius of 20. A pencil beam is positioned at [25, 25, 0], emitting light in the +z direction. |
| | Prompt 3 | I would like to simulate a three-layer structure with the following specifications:<br>• First layer: Extends from z=0 to z=10 with optical properties $\mu_a = 0.01$, $\mu_s = 1.2$, $g = 0.75$, and $n = 1.3$.<br>• Second layer: Extends from z=10 to z=20 with optical properties $\mu_a = 0.04$, $\mu_s = 1.0$, $g = 0.9$, and $n = 1.3$.<br>• Third layer: Extends from z=20 to z=50 with optical properties $\mu_a = 0.15$, $\mu_s = 1.1$, $g = 0.9$, and $n = 1.3$.<br>A planar light source is positioned on the z = 0 plane, spanning the region from [0,0,0] to [40,20,0], and emitting light in the +z direction. |
| | Prompt 4 | I would like to simulate a cubic box spanning from [0 0 0] to [60 60 60] with optical properties $\mu_a = 0.005$, $\mu_s = 10.0$, $g = 0.9$, and $n = 1.3$. A Gaussian beam with a waist radius of 2 is positioned at [30 30 0], emitting light along the direction [0, 0.707, 0.707]. |
| | Prompt 5 | I would like to simulate a cylinder with a radius of 10, a base center at [0 0 0] and a top center at [0 0 30.5]. The cylinder has optical properties $\mu_a = 0.005$, $\mu_s = 0.0$, $g = 0.0$, and $n = 1.4$. A cone-shaped light source is positioned at [0 0 0], emitting light in the +z direction with an emission angle of 5 degrees. |

| Subject ID | Prompt ID | Prompt |
|---|---|---|
| S2 | Prompt 1 | Please prepare an MCX JSON file with the following settings: The whole domain is $90x90x90mm^3$ with a 1 mm edge length voxel size. From a z-position of 0 mm to 30 mm, set the optical properties of the domain to have a $mua = 0.1mm^{-1}$ and a $mus = 0.75mm^{-1}$ and a g = 0.75 and a refractive index (n) of 1.93. From a z-position of 30 mm to 60 mm, set the optical properties of the domain to have a $mua = 0.001mm^{-1}$ and a $mus = 1.05mm^{-1}$ and a g = 0.8 and a refractive index (n) of 1.4. From a z-position of 60 mm to 90 mm, set the optical properties of the domain to have a $mua = 0.004mm^{-1}$ and a $mus = 3.05mm^{-1}$ and a g = 0 and a refractive index (n) of 0.8. Outside of the overall domain, set the properties to air ($mua = 0$, $mus = 0$ , $g = 1$, $n = 1$).<br>Run the simulation with 300,000 photons. The source is a point source centered at [45,0,0] mm propagating in the direction of [0 0 1] mm and simulated over a total simulation time of 5e-8 seconds. Provide the output as volumetric fluence. |
| | Prompt 2 | Please prepare an MCX JSON file with the following settings: The whole domain is $80x90x80mm^3$ with a 1 mm edge length voxel size. From a z-position of 0 mm to 30 mm, set the optical properties of the domain to have a $mua = 0.1mm^-1$ and a $mus = 0.75mm^-1$ and a g = 0.75 and a refractive index (n) of 1.93. From z-position of 30 mm to 80 mm, set optical properties equal to a $mua = 0.001mm^-1$ and a $mus = 1.75mm^-1$ and a $g = 0.75$ and a refractive index (n) of 1.93 . Place a box inclusion at an origin of [30, 30, 50 ] mm of total size [15, 15, 15] mm and optical properties $mua = 0.05mm^-1$ and a $mus = 2.00mm^-1$ and g=0, and n=1.3.<br>Run the simulation with 400,000 photons. The source is a gaussian source centered at [45,45,0] mm propagating in the direction of [0 0 1] mm with a waist radius of 4 mm and simulated over a total simulation time of 5e-8 seconds. Provide the output as output flux. |
| | Prompt 3 | Please prepare an MCX JSON file with the following settings: The whole domain is $100x100x100mm^3$ with a 1 mm edge length voxel size and optical properties of $mua = 0.001mm^-1$ , $mus = 1mm^-1$ , g=0 and n=1.4. Place a cylinder from a starting postition of[50 50 50] mm to [70 70 70] mm with a radius of 4 mm with optical properties of $mua = [0.05]mm^-1$ , $mus = 2mm^-1$, g=-0.8 and n=1.4. The area outside of the domain is set to optical properties of air ($mua = 0$, $mus = 0$ , $g = 1$, $n = 1$).<br>Run the simulation with 1,000,000 photons. The source is a cone source centered at [50,50,30] mm propagating in the direction of [0 0 1] mm with an angle of 6 degrees and simulated over a total simulation time of 5e-8 seconds. Furthermore, a detector of radius 2 mm is placed at [50, 50,100] mm. Provide the output as energy deposited. |
| | Prompt 4 | Please prepare an MCX JSON file with the following settings: The whole domain is $100x100x100mm^3$ with a 1 mm edge length voxel size and optical properties of $mua = 0.001mm^-1$ , $mus = 1mm^-1$ , $g = 0$ and $n = 1.4$. Place a Sphere at a origin position of [50 50 50] mm with a radius of 15 mm with optical properties of $mua = [0.05]mm^-1$ , $mus = 2mm^-1$, $g = -0.8$ and $n = 1.4$. The area outside of the domain is set to optical properties of air ($mua = 0$, $mus = 0$ , $g = 1$, $n = 1$).<br>Run the simulation with 1,000,000 photons. The source is a hyperboloid source centered at [50,50,0] mm propagating in the direction of [0 0 1] mm with a focus of 6 mm and a rayleigh range of 10 mm simulated with a total simulation time of 5e-9 seconds. Furthermore, a detector of radius 6 mm is placed at [50, 50,100] mm. Provide the output as output flux. |

| Subject ID | Prompt ID | Prompt |
| --- | --- | --- |
| | Prompt 5 | Please prepare an MCX JSON file with the following settings: The whole domain is $50x50x50mm^3$ with a 1 mm edge length voxel size and optical properties of $mua = 0.001mm^-1$ , $mus = 1mm^-1$ , $g = 0$ and $n = 1.4$. Place a Sphere at a origin position of [10 10 10] mm with a radius of 5 mm with optical properties of $mua = [0.05]mm^-1$ , $mus = 2mm^-1$, $g = 0.8$ and $n = 1.4$. Place a Sphere at a origin position of [35 35 35] mm with a radius of 10 mm with optical properties of $mua = [0.08]mm^-1$ , $mus = 2.5mm^-1$, $g = 0.8$ and $n = 1.4$. The area outside of the domain is set to optical properties of air ($mua = 0$, $mus = 0$ , $g = 1$, $n = 1$). Run the simulation with 1,000,000 photons. The source is a hyperboloid source centered at [50,50,0] mm propagating in the direction of [0 0 1] mm with a focus of 6 mm and a rayleigh range of 10 mm simulated with a total simulation time of 5e-9 seconds. Furthermore, a detector of radius 6 mm is placed at [25, 25,50] mm. Place another detector of radius 3 mm at [25,25,0] mm. Provide the output as output flux. |
| S3 | Prompt 1 | Can you generate a forward simulation to output the fluence? The domain is a box with x,y,z dimensions (60, 60, 60). The Inside the box is a sphere centered at 25,25,25 with a radius of 10. Please set the sphere coefficients, mua to be 0.15, mus to be 1.1, g is 0.9 and n is 1.3. Please also set the z layers 0 through 20 of the box to have the following coefficients, mua is 0.01, mus is 1.2, g is 0.75, n is 1.5. The source is a pencil beam with direction (0,0,1) at position (25,25,0), launching 100000 photons. Thank you |
| | Prompt 2 | Can you please run a forward simulation that will output the energy deposit. The domain is made up of a cylinder with one circular end located at (5,5,5) and the other circle located at (5,5,30. The cylinder has a radius of 2. The mua is 0.2, mus is 1.5, g is 0.85 and n is 1.45. The cylinder is located in a grid box with dimensions (60, 60, 60), with coefficients mua and mus are 0 and g and n are 1. The source is isotropic located at (15, 15,0) with a direction of (0,0,1). |
| | Prompt 3 | I need to obtain output flux from a forward simulation using a gaussian source positioned at (15,15,15) with a direction of (0,1,1) and a waist radius of 1.5). The domain is box with an origin of (10,10,10) and a size of (5,5,10). The box ahs optical properties mua is 0.1, nus is 1.1, g is 0.88 And n is 1.37. Inside this box is a sphere centered at (25, 25,25) with a radius of 5. Its optical properties are mua is 0.2, mus is 0.85 g is 0.9 and n is 1.33. Background optical properties are mua and mus are 0 and g and n are 1. |
| | Prompt 4 | Please run a simulation that uses a source of 600000 photons cone source positioned at (30,15,15) with a direction of (-1,0,0). Inside the domain there is a box with dimensions (6 6 6) positioned at (8,8,8). The box has optical properties, mua is 0.2 mus is 0.75, g is 0.88 and n is 1.36. Please set the z layers 0 to 25 to have mua of 0.1, mus of 0.8, g is 0.9 and n is 1.33. I need the fluence as the output of the simulation. |
| | Prompt 5 | Can you please generate a forward MCX simulation to output the fluence? The domain is a grid box with x, y, z dimensions of (70, 70, 70). Inside the box is a sphere centered at (35, 35, 30) with a radius of 8. Please set the sphere optical properties to mua = 0.18, mus = 1.0, g = 0.9, and n = 1.35. Please also set the z layers 0 through 18 of the box to have optical properties mua = 0.02, mus = 1.25, g = 0.8, and n = 1.45. The source is a pencil beam positioned at (35, 35, 0) with a direction of (0, 0, 1), launching 200000 photons. Thank you. |

| Subject ID | Prompt ID | Prompt |
|---|---|---|
| S4 | Prompt 1 | create an mcx simulation of a two layered domain of size 100x50x60 voxels, with the top z-layer extending between z=1 and 10 filled with medium of mua=0.1/mm, mus=1/mm, g=0.01 and n=1.37; the second z-layer extend from 11 to 60, filled with mua=0.02/mm, mus=0.3/mm, g=0.9, and n=1.4; place a disk source of radius 2 at [50, 30, 0] with an incident direction of [0, 0, 1]. Simulate 1e6 photons with a maximum time gate length of 5e-9 second, and a time step of 1e-10 second |
| | Prompt 2 | simulate two spheres. one centered at [30, 10, 20] with radius of 10, another one centered at [30, 50, 30] with a radius of 5, inside a 80x80x40 voxelated grid. The first sphere has mua=0.003/mm, mus=0.8/mm, g=0, and n=1.34; the second sphere has mua=0.01/mm, mus=0.1/mm, g=0.89, n=1.37; the background medium is filled by mua=0, mus=0.0001/mm, g=0.9, n=1. Place a pencil source at [30, 0, 10], with an incident direction of [0, 1, 0]. Simulate 1000000 photons and output fluence rate. |
| | Prompt 3 | create a JSON input file for an MCX simulation over a domain of size 100x100x100, the domain contains a spherical inclusion with center located at [50, 50, 50] and a radius of 30; the domain also contains a cylinder with two ends located at [50, 50, 0] and [50, 50, 100], with a radius of 10; the sphere is filled with medium with absorption coefficient (mua) of 0.07/mm, scattering coefficient (mus) of 1/mm, anisotropy (g) of 0.89, and refractive index (n) of 1.3; the cylinder has mua=0.001/mm, mus=0.002/mm, g=0.8, and n=1.4; the background medium of the domain has mua=0, mus=0, g=1, and n=1. The source is located at [50, 40, 0], launching photon at direction [0, 0, 1]. Please simulate 10000 photon packets and return the fluence output for a maximum time window of 1e-9 second. |
| | Prompt 4 | Create a simulation for a 60x50x40 voxelated space filled with scattering medium with absorption coefficient (mua) of 0.01/mm, scattering coefficient (mus) of 5/mm, anisotropy (g) of 0.9, and refractive index (n) of 1.37; place a disk source of radius of 2 at position [30, 20, 1]; add a zlayer filled with medium type 0 at z=1; run 100000 photon packets and save diffuse reluctance output. |
| | Prompt 5 | Help me build an MCX input file for a simulation inside a 100x80x50 voxelated space with a voxel size of 0.5mm. A pencil beam source is placed at the center on the top face of the domain, pointing downwards. The space is filled with mua=0.002 per mm, mus=0.7 per mm, g=0.1, and refractive index n=1.3. Create 3 random spherical objects inside the space, with random center positions and radii, filled with random optical properties. Two detectors are located at [50, 30, 50] and [50, 40, 50], both with an radius of 2. Please run 1e7 photons and save detected photon data and the fluence. |
